# Transposable elements orchestrate subgenome-convergent and -divergent transcription in common wheat

**DOI:** 10.1101/2022.10.14.512211

**Authors:** Yuyun Zhang, Zijuan Li, Jinyi Liu, Yu’e Zhang, Luhuan Ye, Yuan Peng, Haoyu Wang, Huishan Diao, Yu Ma, Meiyue Wang, Yilin Xie, Tengfei Tang, Yili Zhuang, Wan Teng, Yiping Tong, Wenli Zhang, Zhaobo Lang, Yongbiao Xue, Yijing Zhang

## Abstract

The success of common wheat as the global staple crop is derived from genome diversity and redundancy as a result of allopolyploidization [1-3], giving rise to the major question how the divergent and convergent transcription among different subgenomes are achieved and harmonized in a single cell. The regulatory information is largely encoded in DNA regulatory elements (REs) interpreted by sequence specific transcription factors (TFs). Here, we created a catalog of genome-wide TF-binding sites (TFBS) to assemble an extensive wheat regulatory network comprising connections among 189 TFs and 3,714,431 REs, which enhances the understanding of wheat regulatory mechanisms on an unprecedented scale. A significant fraction of subgenome-divergent TFBS are derived from recent subgenome-asymmetric expansion of particular transposable element (TE) families. In contrary, TFBS derived from ancient TE expansion largely underwent parallel purifying selection during independent evolution of each subgenome, despite extensive unbalanced turnover of flanking TEs. Altogether, the subgenome-convergent and -divergent regulation in common wheat is orchestrated via differential evolutionary selection on paleo- and neo-TEs.

## Introduction

Polyploidy is one major factor driving plant evolution and speciation, which is particularly prevalent in plants[1-4]. Indeed, all angiosperms have undergone paleopolyploidy events, and a significant proportion of crops have experienced polyploidy in more recent times [5]. Allopolyploids potentially benefit from fixed “hybrid vigor”[4, 6], showing higher adaptability and plasticity compared to the progenitors. This is attributed to diversity and synergy among different subgenomes[7, 8], raising the major question about how the subgenome-divergent and -convergent regulation are achieved and harmonized in polyploids.

Common wheat (*Triticum aestivum*) converged three subgenomes and established as the global staple crop. The gene composition is largely conserved across subgenomes, whereas the intergenic sequences are almost completely turned over [9, 10]. There are abundant regulatory elements (REs) present in intergenic regions in wheat [11-13], the variation of which affect a wide range of agronomic traits [14-17]. Intergenic RE decay may help explain the finding that 30% of wheat homoeologs showed nonbalanced expression [12, 13, 18], while it is still not clear how the RE divergence across subgenomes are specifically interpreted to dictate subgenome diversified transcription. Furthermore, despite the highly diversified regulatory regions, extensive coordination of homoeologs was detected throughout development [18, 19], giving rise to a more tangled question how this evolutionary constraint on transcriptional regulation is achieved. Specific recognition and binding of REs by transcription factors (TFs) are the primary mechanism by which cells interpret genomic features and regulate genes. Elucidating the extent to which TF binding differences occur across subgenomes, and the global relationship between TF binding and subgenome variation of REs is the key to address the above questions.

## Results

### Genome-wide profiling of TFBS in common wheat

We cloned 189 TFs from 30 families, including 107 highly expressed TFs and 82 TFs whose functions were reported or were among the hub TFs of the co-expression network (Table S1). Each clone was verified by full-length cDNA sequencing to confirm a lack of chimeric fragments from homologs. Next, DAP-seq [20] was performed to characterize the genome-wide binding of these TFs, which were classified according to the *de novo* identification or enrichment of canonical representative motifs, resulting in 45 high-, 48 median-, and 97 low-confidence TF datasets (HC, MC and LC) (Fig. 1a and Fig. S1). The HC and MC were used for subsequent analysis. Given that noncanonical binding may also be functional (Fig. S2) [21, 22], all DAP-seq data, including LC TFs, and peak files were deposited in a public database (GSE192815). The DAP-seq success rate varied among TF families, AP2, MYB and B3 TF families displayed high, median and low success rates, respectively (Fig. 1b). The binding for the TF families with low success rates likely require co-factors. All data were visualized through a customized genome browser (http://bioinfo.sibs.ac.cn/dap-seq_CS_jbrowse/). The TFBS for the AP2 TFs were grouped together more than the TFBS for the other TF families (Fig. 1c), indicating the evolutionary constraint of AP2 functions. The binding specificities of homologous TFs were generally conserved across subgenomes (enlarged heatmap in Fig. 1c), implying that the subgenome divergent regulation likely embedded in subgenome divergent REs.

**Figure 1.**
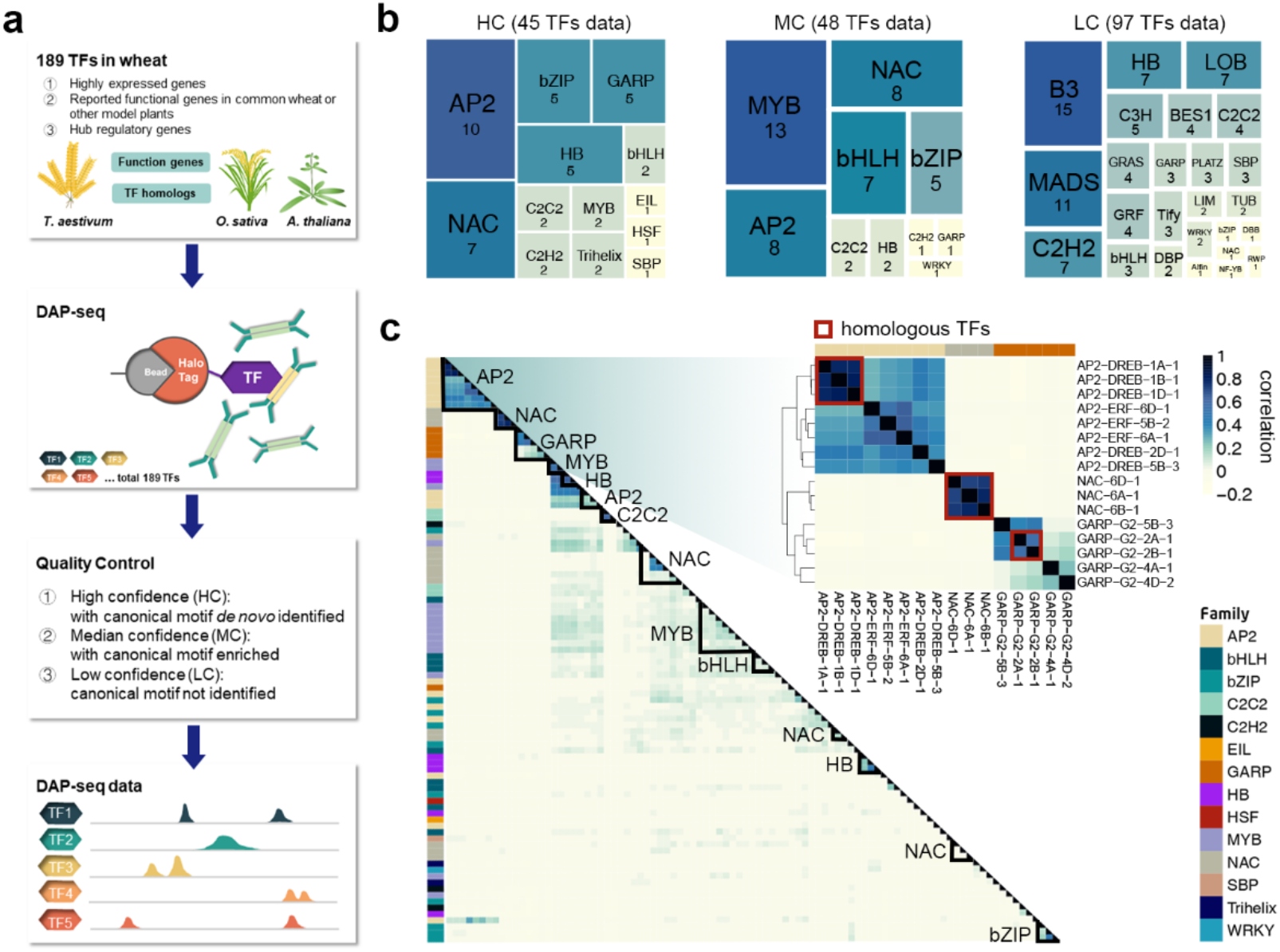
Genome-wide binding of wheat transcription factors. **a**, Schematic of the experimental design and filtering steps. **b**, Tree map illustrating the fraction of TF families with high-confidence (HC; with canonical motif de novo identified), median-confidence (MC; with canonical motif enriched), and low-confidence (LC; canonical motif not enriched) TFs. **c**, Clustering of TF binding correlations according to the occurrence of DAP-seq peaks indicating that TFs from the same family and the same orthologous gene group generally have similar binding profiles. The heatmap of the clustering of AP2, NAC, and GARP family TF binding correlations is enlarged on the right; colored boxes represent the homologous TFs in the heatmap.

The TFBS are not randomly distributed throughout the genome, with highly-occupied clusters with 42,332 binding events designated as high-occupancy target (HOT) regions [23, 24]. The high regulatory activities of HOT regions were reflected by the relatively high levels of chromatin openness, epigenetic activity, and conservation across wheat species (Fig. 2a). Additionally, 80% of the HOT regions had highly conserved sequences among subgenomes (Fig. 2b), mostly localized to gene-proximal regions (Fig. 2c). By comparing HOT regions with higher-order chromatin structures, we determined that HOT regions were preferentially localized to TAD boundaries (Fig. 2d). Figure 2e presents the genomic features of one subgenome-conserved HOT region. Although the local chromatin structure varied substantially across subgenomes, HOT regions were still preferentially localized to TAD boundaries. Previous research indicated TADs are formed via promoter-enhancer linkages mediated via coopted regulatory elements [25]. Here, the considerable enrichment of HOT regions in TAD boundaries implies that a high TF occupancy may facilitate TAD formation.

**Figure 2.**
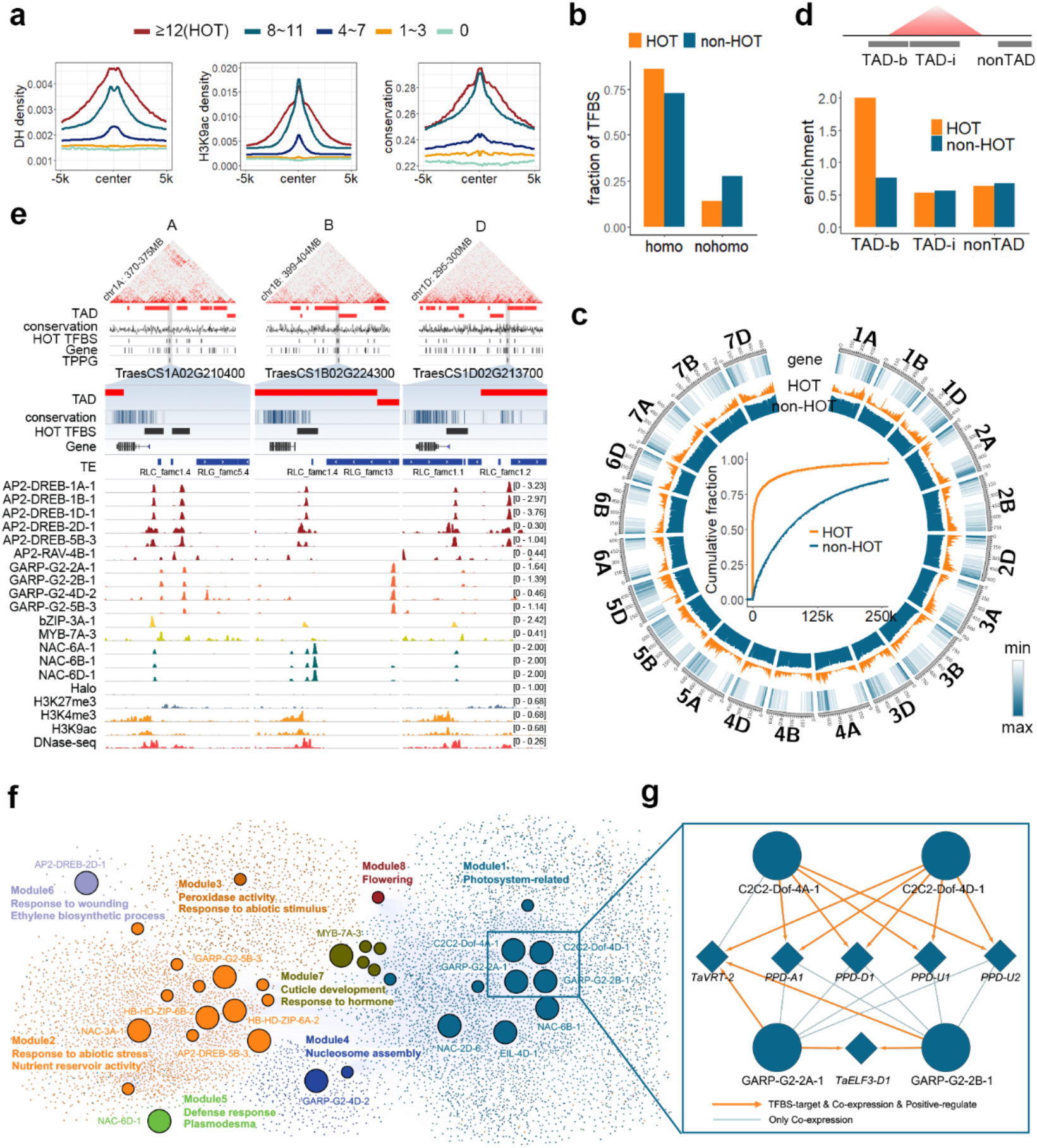
Chromosomal features of regulatory circuitry. **a**, DHS and H3K9ac read density and conservation of TFBS, which were grouped according to the number of bound TFs. Figures present the average signal densities and conservation score at 100-bp resolution within a 10-kb window centered on merged TFBS centers. Merged TFBS with more than 12 binding sites were identified as HOT regions. **b**, Fractions of HOT regions and non-HOT regions overlapping with subgenome-homologous and -nonhomologous regions. **c**, Outer: Circos plot presenting the genomic distribution of HOT and non-HOT regions. Inner: cumulative fractions of the distance between a gene TSS and HOT regions or non-HOT regions. **d**, Enrichment of HOT regions and non-HOT regions in the TAD boundary, TAD internal regions, and non-TAD regions. **e**, Genomic tracks illustrating the targeting of the 1:1:1 homologous gene *TPPG* by a subset of TFs. The *TPPG* promoters are located in subgenome-homologous HOT regions and the TAD boundary. **f**, Regulatory circuit integrating information from TF–target pairs and coexpression modules. Node colors represent different coexpression modules. Circle and diamond nodes represent high- or median-confidence TFs and non-TFs, respectively. **g**, Enlarged image of nodes. Lines for different connection types are indicated below.

TFs guide cellular development and activities through the highly cooperative and dynamic control of gene expression. By integrating analyses of TF binding and coexpression profiles for diverse tissues (see Methods), we built a regulatory network comprising 8 modules, with connections among 34 TFs and 8,937 genes. To characterize these modules, we screened for enriched Gene Ontology terms using GOMAP [26]. The functionally annotated groups are summarized in Figure 2f and Figure S3. A module comprising TFs and targeted genes potentially involved in photosynthesis is presented in Figure 2g (zoomed in on the right), comprising the well reported TFs and factors involved in photoperiod and photosynthesis, including Dof, Ppd1 and Elf3[27]. This information allowed us to comprehensively explore the evolution of polyploid regulation and the underlying mechanism.

### Subgenome A specific expansion of RLG_famc7.3 contributed to subgenome A biased TF binding

To compare the regulatory architecture across subgenomes, HC TFBS were divided according to their sequence conservation (Fig. 3a). On average, 40% of the TFBS were localized in nonhomologous regions across subgenomes (i.e., pervasive asymmetric subgenome regulation). To examine the different functions among genes regulated by subgenome-convergent and -divergent TFBS, we searched for the over-represented GOMAP terms associated with genes preferentially containing nonhomologous and homologous TFBS, respectively. Genes regulated by homologous TFBS were mostly involved in relatively conserved biological pathways including membrane architecture and development (Fig. 3b), whereas genes associated with subgenome-divergent regulation were mostly involved in defenses. Thus, subgenome-divergent environmental adaptation may be encoded by the subgenome-divergent regulatory circuit.

**Figure 3.**
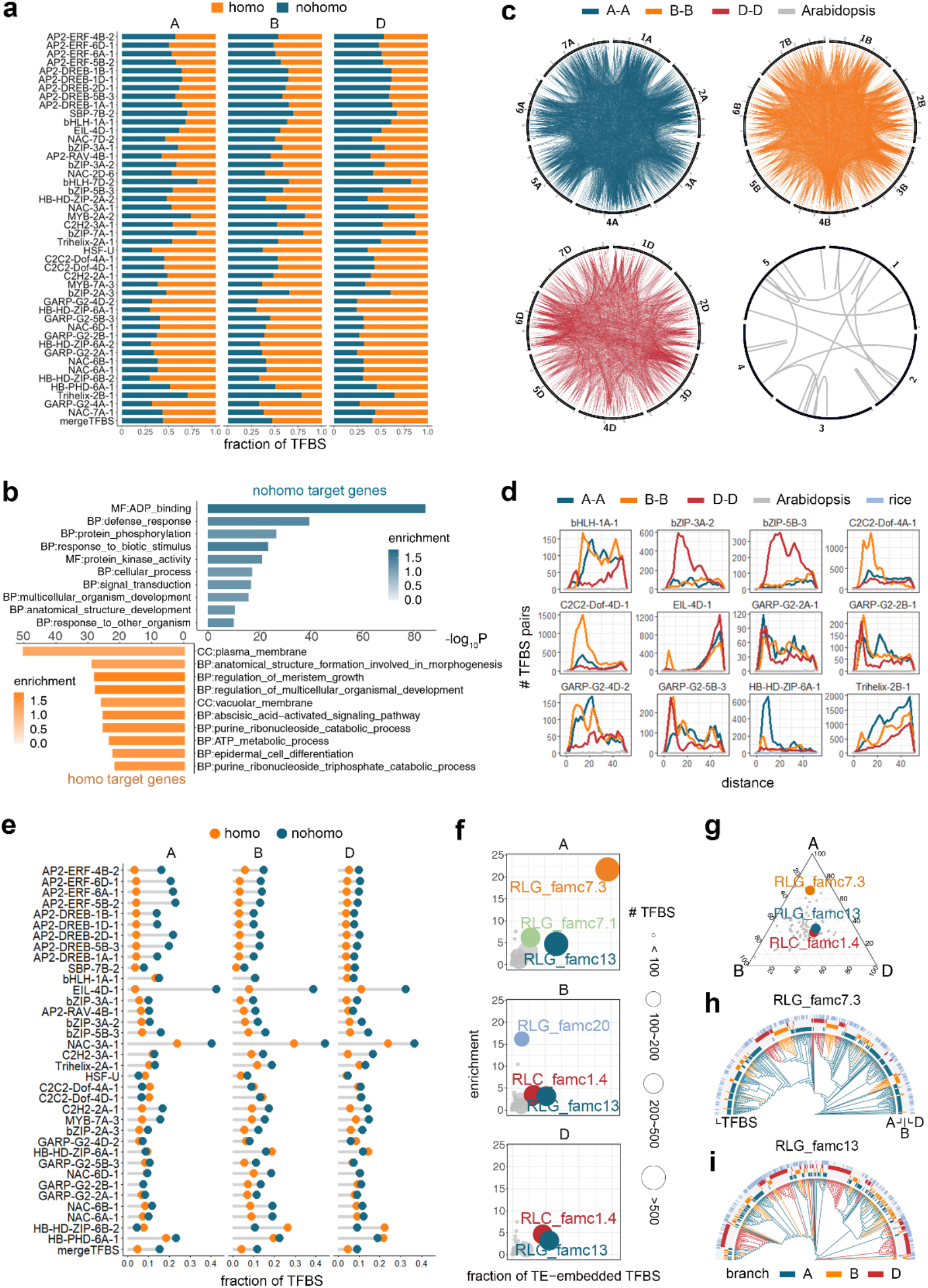
Subgenome A-specific expansion of RLG_famc7.3 contributed to subgenome A-biased TF binding. **a**, Fraction of subgenome-homologous and -nonhomologous TFBS. **b**, Top enriched GO terms for homologous and nonhomologous TFBS-biased targeted genes ranked according to the enrichment *P* value. **c**, Circos plots showing the bHLH-1A-1 TFBS pairs with high sequence similarity within each subgenome; the homolog bHLH28 in *Arabidopsis thaliana* was shown as control. For each TF peak set, 1,500 peaks were randomly selected. The calculation of sequence similarity is described in Methods. **d**, Sequence distance distribution between TFBS in each subgenome of CS and TFBS of homologous TFs in *Arabidopsis thaliana* (grey line) and *Oryza sativa* (light blue line). **e**, Fraction of subgenome-homologous and -nonhomologous TFBS embedded in TEs within open chromatin regions. **f**, Enriched TE subfamilies harboring nonhomologous TFBS. **g**, Relative abundance of TE subfamilies across subgenomes. The top enriched TEs in Fig. 2f are highlighted. **h**, Dendrogram presenting the sequence similarity between full-length RLG_famc7.3 members. **i**, Dendrogram presenting the sequence similarity between full-length RLG_famc13 members.

We next asked the origin of subgenome divergent TF binding. For each TF, subgneome nonhomologous TFBS were collected whose pair-wise sequence similarity were calculated within each subgenome. The circos plot in Fig. 3c connected bHLH-1A-1 binding sites showing high pair-wise sequence similarity in each subgenome, which is much more abundant as compared to that in Arabidopsis. The distribution of pair-wise TFBS sequence distance for all TFs were shown in Fig. 3d and Fig. S4. It is clear that almost all nonhomologous TFBS underwent one or multiple round(s) of expansion event in each subgenome, as reflected by the apparent peak(s) indicating sequence similarity among a number of TFBS, which were not happened in other well-studied model plants including Arabidopsis and rice (Fig. 3c-d and Fig. S4). Further examination revealed that majority of (>80%) TFBS showing high pair-wise sequence similarity are embedded in TEs (Fig. S5), whereas the fraction is lower than 40% for other TFBS without high sequence similairy. This is as expected given that more than 85% of the wheat genome consists of TEs, expansion of which with built-in regulatory copies may quickly rewire transcriptional patterns and lead to novel functions and increasing regulatory complexity [28-32]. By calculating the proportion of nonconserved regions in TEs, we determined that 80%–90% of the evolutionary variance among subgenomes can be explained by TE domestication (Fig. S6), 30% of which were localized to open chromatin regions, representing highly active binding sites *in vivo* (Fig. 3e). Thus, it is likely that TEs contributed a vast TF occupancy difference across subgenomes.

Although TFBS overlapped many TE types, significant proportions of subgenome A-specific binding involved RLG_famc7.3, whereas RLG_famc13 contributed to the nonconserved binding in all three subgenomes (Fig. 3f). The enrichment of different TE families in subgenome-specific TFBS indicates the high plasticity of the regulatory framework was shaped by TE-embedded TFBS. To trace the expansion history, full-length RLG_famc7.3 and RLG_famc13 members were used to construct an evolutionary tree. Both TE families expanded after the subgenomes divergence. RLG_famc7.3 specifically expanded in subgenome A (Fig. 3g), 69% of which present in subgenome A (Fig. 3h). Whereas RLG_famc13 expanded in all three subgenomes following the divergence from common ancestors (Fig. 3i). The TEs with TFBS were dispersed in the TE subfamily clusters, reflecting their diverse origins (Fig. 3h-i). Altogether, recent subgenome asymmetric expansion of particular TE families is an important source of subgenome divergent TFBS.

### Impact of ancient TE exaptation on subgenome balanced transcription

To characterize the regulatory consequences of subgenome balanced and unbalanced TF binding, we focused on gene-proximal TFBS. Recent transcriptomic data indicated that at least 30% of subgenome conserved triad genes exhibited unbalanced expression [18], which is likely coordinated by RE sequence context, epigenetic modifications, and TF regulation [12]. To clarify this divergence, we quantitatively partitioned TFBS in triad promoters according to subgenome-preferential binding (Fig. 4a and Methods). The AP2 occupancy profile was stable across subgenomes, whereas the GARP and NAC bindings are highly diverse across subgenomes (Fig. 4b–c). The subgenome-unbalanced binding of triads was consistent with their subgenome-unbalanced expression (Fig. 4d).

**Figure 4.**
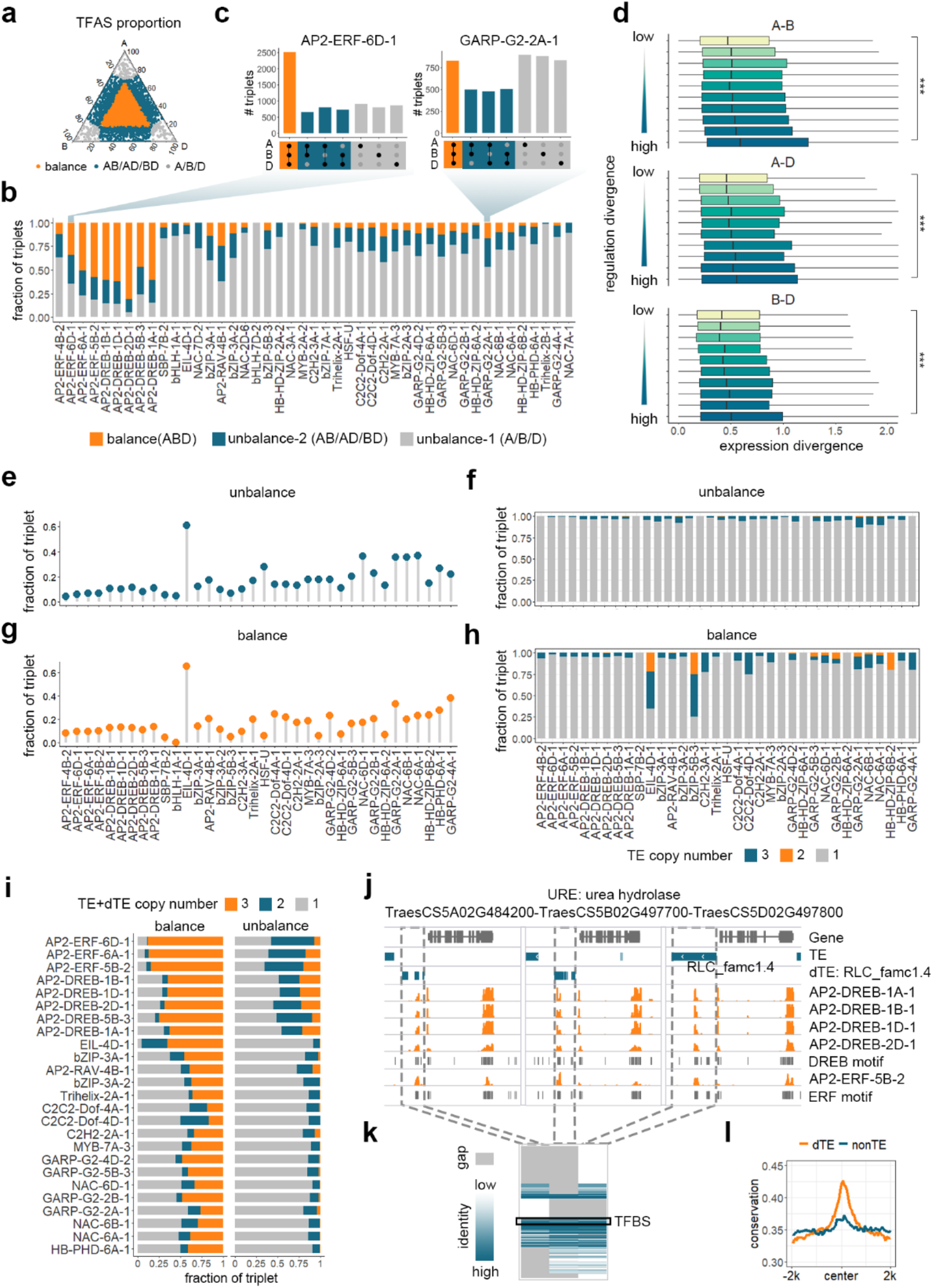
Subgenome parallel evolution of TE-derived TFBS contributed to subgenome-convergent transcription. **a**, The quantitative divergence of AP2-ERF-6D-1 binding in the promoter of triad genes. Orange dots represent the balance regulatory pattern. Blue and grey dos represent the unbalance regulatory pattern. **b**, Fraction of balanced and unbalanced TF binding in the promoters of triad genes (1:1:1 correspondence across subgenomes). **c**, Bar plot presenting the results of the quantitative analysis of the balanced and unbalanced TF binding of AP2-ERF-6D-1 and NAC-6A-1. **d**, Subgenome divergent expression of different groups of orthologous gene pairs. Gene pairs were grouped according to the regulatory divergence, which was calculated as the sum of the binding of all TFs |log_2_fc|. The expression divergence was calculated as the |log_2_fc| of two orthologous genes. The Wilcoxon signed-rank test was used to compare the expression divergence of different groups. \*\*\**P* < 0.001 (H1: the expression divergence of the gene pairs with the lowest regulatory divergence differed from that of the gene pairs with the highest regulatory divergence). **e**, Fraction of unbalanced triads targeted by TE-derived TFBS. **f**, Fraction of TE copy numbers in the promoters of unbalanced triads. **g**, Fraction of balanced triads targeted by TE-derived TFBS. **h**, Fraction of TE copy numbers in the promoters of balanced triads. **i**, Fraction of TE and degenerated TE (dTE) copy numbers of balanced and unbalanced triads targeted by TE-derived TFBS. **j**, Genome tracks illustrating the dTEs contributing to the convergent TF binding of triad gene *URE* promoters. **k**, Multiple sequence alignment of the TEs and dTEs in Fig. 3f. The black box indicates the TFBS. Blue and gray represent the aligned nucleotides and alignment gaps, respectively. The color scale reflects the alignment identity. **l**, Conservation score profile of dTE-derived TFBS and non-TE TFBS. The conservation score (bin size 50 bp) within a 4-kb window centered on merged TFBS centers is provided. TFBS in triad promoters without a TE in all three subgenomes were considered as non-TE TFBS. Conservation score is a measure of sequence conservation across wheat species.

We further evaluated the relationships between TE and subgenome unbalanced binding in triad promoters. 10%–40% of unbalanced binding was explained by asymmetric TE insertions (Fig. 4e). For triads with TE-embedded TFBS in at least one promoter, more than 90% of the unbalanced TF binding was due to TEs inserted in only one subgenome (Fig. 4f), indicating unbalanced TE insertion is one major source creating subgenome-unbalanced TF binding.

Surprisingly, for TFs with subgenome-balanced binding across triads, similar fractions were contributed by TEs (Fig. 4g). Moreover, most TEs contributed to TFBS (>95%) were detected in only one subgenome (Fig. 4h). To elucidate the mechanism underlying the balanced TF occupancy when the contributing TEs are unbalanced across triad promoters, we restricted our analysis primarily to the triads with at least one member with TE-embedded TFBS in its promoter. Because of the long evolutionary history of TEs in wheat, there are many TE relics in the genome, which must be considered when characterizing the dynamic effect of TE proliferations[9, 32]. We detected the degenerated TEs (dTEs) in triplet promoters (Methods) and analyzed their overlap with TFBS (Fig. S7). When TEs and dTEs were considered together, a much higher fraction of balanced triplets were targeted by TE- or dTE-derived TFBS in all three triad members (Fig. 4i, left panel), and the fraction is up to 89% for AP2 families; whereas for the unbalanced triplets, the contribution of dTEs to the TFBS was rather limited (~7% on average) (Fig. 4i, right panel). The biological significance of these dTE-derived TFBS is supported by the high conservation across wheat species with different ploidy levels; in contrast, the flanking TE sequences are highly diverse (Fig. 4j-l). This is an intriguing finding, indicating that a significant fraction of TFBS derived from anciently expanded TEs experienced parallel purifying evolution for each individual subgenome after divergence, whereas the flanking TE sequences underwent relaxed selection or diversifying selection, resulting in subgenome-unbalanced decay. Despite we do not expect that all these TE domestication events have functional implications, the specific evolutionary constraint on TE-derived TFBS and the association between balanced binding and balanced expression, provide important evidence of the evolutionary effects of ancient TE remnants on subgenome-convergent transcriptional regulation.

### Preferential contribution of paleo-expansion of RLC_famc1.4 to TE-derived subgenome convergent TF binding

Both RLC_famc1.4 and degenerated RLC_famc1.4 are the top ranked TE families contributed to balanced TFBS across triad promoters (Fig. 5a), accounting for a total of 23% balanced TE-derived TFBS targets. It is interesting that the ancient expansion of almost all TF families profiled here are preferentially contributed by RLC_famc1.4 (Fig. 5a). Why the specific TE family dominate the TFBS exaptation in gene proximal region is an interesting issue. Unlike subgenome specific expansion of RLG_famc7.3 and RLG_famc13, a mixture of RLC_famc1.4 TEs from three subfamilies were observed in the phylogenetic tree (Fig. 5b), indicating most RLC_famc1.4 TEs may derived from the common ancestor. In support of this, the age distribution of transposition events suggested that RLC_famc1.4 members are relatively ancient among the genome-wide abundant TEs (Fig. 5c), as indicated by the low similarity between left and right LTR regions [33]. To further trace the RLC_famc1.4 expansion events, we compared the pair-wise sequence similarity for full-length RLC_famc1.4 TEs, which revealed one round of RLC_famc1.4 expansion predated the divergence between wheat and rye (Fig. S8). Another new expansion event occurred in rye, whereas no apparent RLC_famc1.4 dispersion was detected in barley. Thus, both ancient and recent RLC_famc1.4 explosive expansions were observed in Triticeae, which have been significantly and continuously shaping regulatory circuitry related to Triticeae genome evolution and adaptation.

**Figure 5.**
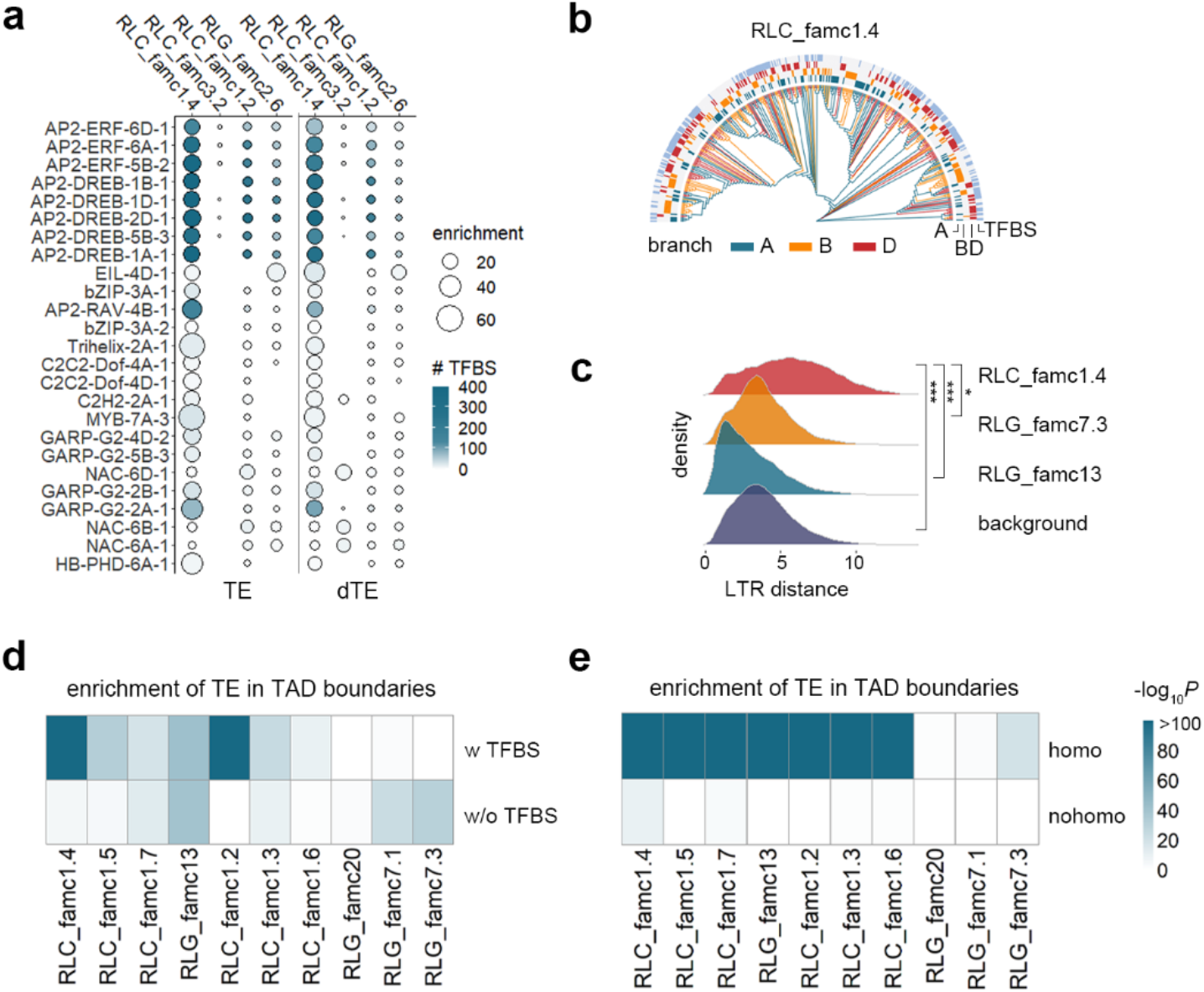
Ancestral expansion of RLC_famc1.4 dominates TE-derived subgenome convergent TF binding. **a**, Enrichment of TE (left) and dTE (right) families contributing to the balanced TF binding across triad promoters. **b**, Dendrogram presenting the sequence similarity between full-length RLC_famc1.4 members. **c**, Age distribution of TEs enriched in TFBS as determined according to the sequence similarity of the LTR at both ends. The Wilcoxon signed-rank test was used to compare the LTR distance of different groups. *0.01 < *P* < 0.05, \*\*\**P* < 0.001 (H1: the LTR sequence of a TE subfamily differed from all TE backgrounds). **d**, Enrichment of TE subfamilies in TAD boundaries with and without TFBS. **e**, Enrichment of TE subfamilies in subgenome-homologous and -nonhomologous TAD boundaries.

## Discussion

In summary, these cistrome maps in common wheat provide a valuable resource to evaluate the integrated interaction of cis- and trans-factors in determining regulatory specificity. We detected different evolutionary forces act on the paleo- and neo-TE-derived TFBS, which orchestrate subgenome-divergent and convergent TF binding, imposing distinctive and synergistic regulatory consequences on polyploidy evolution (Fig. 6).

**Figure 6.**
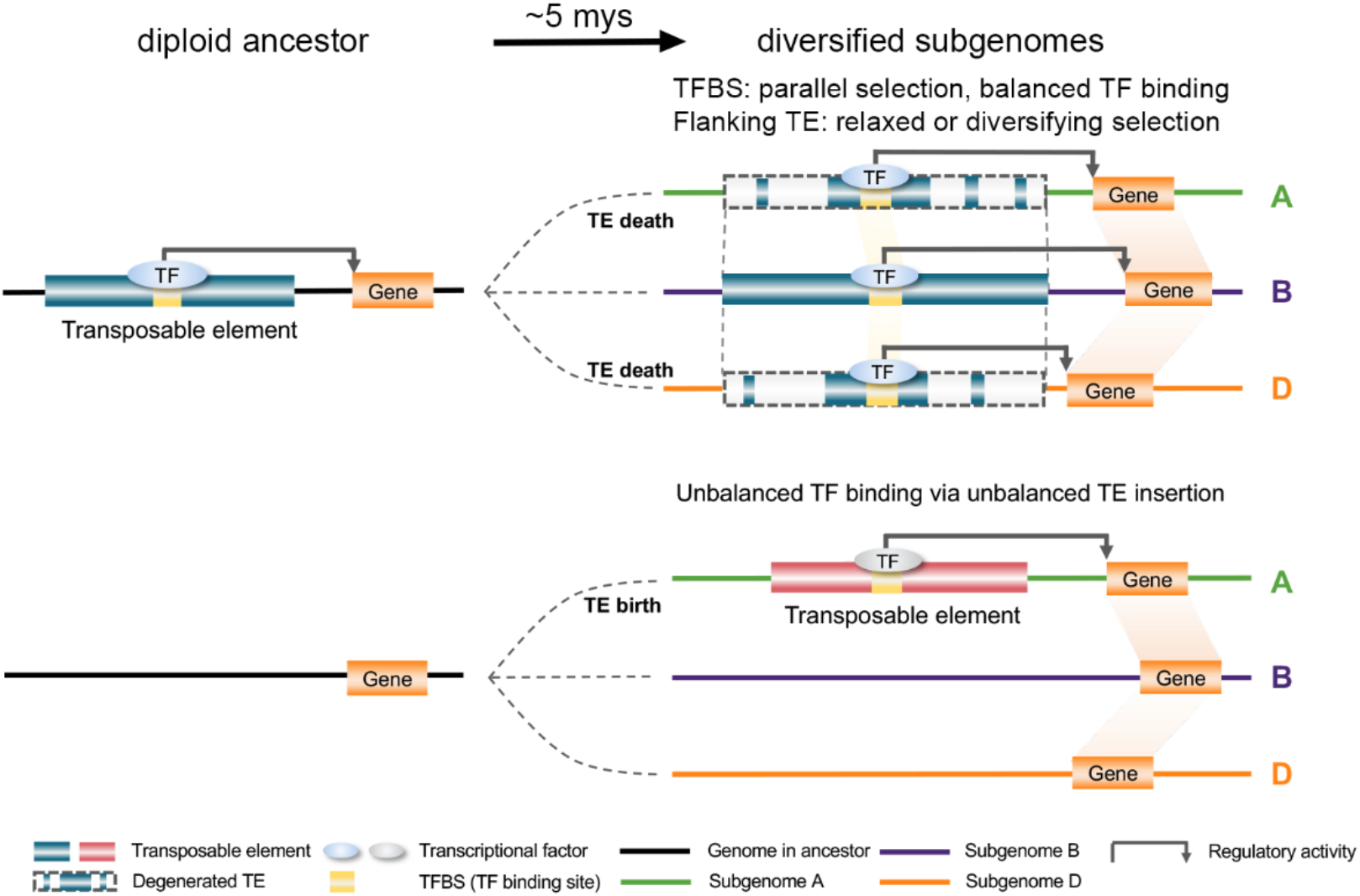
Model illustrating the impact of TE-derived regulatory conservation and innovation on subgenome-convergent and -divergent TF binding and transcription. Top: TFBS embedded in TEs expanded as TEs spread in common ancestors. In most cases, these TFBS were maintained via parallel purifying selection during independent evolution of each subgenome after divergence, whereas the flanking TE sequences degenerated to varying extents (TE death), resulting in subgenome-balanced TFBS and subgenome-unbalanced TEs. This balanced TF binding is associated with subgenome-balanced transcription. Bottom: TFBS embedded in TEs underwent subgenome-specific expansion after divergence (TE birth), which was responsible for subgenome-divergent TF binding and diversified transcriptional regulation.

Rounds of wheat-specific TFBS expansion were detected, which cannot be observed in other model plants including Arabidopsis and rice (Fig. 3c-d). This could be attributed to the active expansion of retro-elements with built-in TFBS in wheat. The TE-embedded TFBS expansion events predate and after subgenome divergence contributed to subgenome convergent and divergent TF regulation, respectively, reflecting the essential role of TE domestication on subgenome regulatory conservation and innovation. The TE exaptation preferentially restricted to a limited number of TE families; RLC_famc1.4 expansion in common ancestors is responsible for subgenome balanced regulation (Fig. 4), and subgenome unique expansion of RLG_famc7.3 resulted in subgenome divergent regulation (Fig. 3). This is analogous to the human specific dispersion of Alu elements, which participated in a variety of human specific regulatory events including conferring enhancer elements and modulating higher-order chromatin structure [34]. Altogether, TEs significantly and continuously rewire wheat regulatory circuit, which also reflected the high plasticity of transcriptional regulation in common wheat.

Despite major changes in REs across subgenomes by TE turnovers, the overall regulatory architecture was highly conserved, both in the convergent evolution of TE-derived TFBS and the extensive coordination of homoeologs (Fig. 4). Since McClintock first described TE functionalization, there has been mounting evidence regarding the profound functional implications of TEs for the regulatory network in animals [29, 35, 36] and plants[28, 37, 38]. However, TEs are subjected to rapid turnover, and the regulatory roles of TEs were mostly associated with creating new TFBS. The TE relics and their evolutionary and functional importance regarding regulatory evolution are unclear, but they are crucial for deciphering the evolutionary impact of TEs on the genome-scale regulatory circuit. As a young polyploid with highly plastic genomes shaped by abundant TEs of various ages, common wheat is ideally suited for subgenome comparisons aimed at clarifying the progressive and ongoing role of TE expansion and degeneration on regulatory evolution. A recent genome-wide characterization of common wheat TEs revealed the evolutionary constraints on the relative distance of specific TE families to genes, a pattern that is conserved across subgenomes [9], suggesting that some TE families may be evolutionarily selected in subgenome ancestor in favor of regulating adjacent genes. In support of this, the present study demonstrated that position-specific selection is a hallmark of the evolution of ancient TE expansion-derived TFBS, and identified specific TE families responsible for subgenome common TF binding.

It is still a mystery why these TFBSs are convergently and specifically selected in different subgenomes after diverged from their common ancestor, and why RLC_famc1.4 dominates the contribution. This may be explained by the recently proposed theoretical model ‘enhancer runaway’ [39, 40], in which selection pressure acts on enhancers to increase gene expression levels, and enhancers that contribute more to protein production are particularly favored. In addition, recent genetic evidence from human study indicated that TEs contributed to chromatin looping and gene regulation [41, 42]. By overlapping with chromatin local structure, we observed that RLC_famc1.4, particularly those overlapping TFBS, were top enriched in TAD boundaries among the genome abundant TEs (Fig. 5d). This enrichment is apparently only for subgenome-homologous TADs (Fig. 5e), indicating the parallel purifying selection on TE-derived TFBS may be associated with subgenome-convergent local chromatin structure. Altogether, these findings demonstrated how different subgenomes of common wheat achieve regulatory harmony from highly diversified regulatory system.

## Methods

### Plant materials and growth conditions

Common wheat (*Triticum aestivum* cultivar “Chinese Spring”) seeds were surface-sterilized via a 10-min incubation in 30% H_2_O_2_ and then thoroughly washed five times with distilled water. The seeds were germinated in water for 3 days at 22 °C, after which the germinated seeds with residual endosperm were transferred to soil. The seedlings (above-ground parts) were harvested after a 9-day incubation under long-day conditions. The harvested samples were frozen in liquid nitrogen for DAP-seq assay.

### DAP-seq assay

DAP-seq was performed as previously described [20]. Genomic DNA was extracted from wheat leaves using Plant DNAzol Reagent (invitrogen) and fragmented. DNA was then end repaired using the End-It kit (Lucigen) and A-tailed using Klenow (3′–5′ exo-; NEB). Truncated Illumina Y-adapter was ligated to DNA using T4 DNA Ligase (Promega). Full length TF was cloned into pIX-Halo vector. Halo-tagged TF was expressed *in vitro* using TNT SP6 Coupled Wheat Germ Extract System (Promega). Halo-TF was immobilized by Magne HaloTag Beads (Promega) and then incubated with the DNA library. TF specific binding DNA was eluted for 10 min at 98°C and amplified with indexed Illumina primer using Phanta Max Super-Fidelity DNA Polymerase (Vazyme). Meanwhile, to capture background DNA which captured by Halo, pIX-Halo vector without TF cloned was expressed and incubated with the DNA library as well. The PCR product was purified using VAHTS DNA Clean Beads (Vazyme) and then sequenced by Novogene (Beijing, China) with the Illumina NovaSeq 6000 system to produce 150-bp paired-end reads.

### Processing of DAP-seq, ChIP-seq, RNA-seq and DHS data

We downloaded the histone ChIP-seq data of 7 typical tissues and 8 external stimuli, seedling RNA-seq data and seedling DNase-seq data of Chinese Spring (CS) from NCBI GEO database (accession number GSE139019 and GSE121903) [11, 12], *OsHOX24* ChIP-seq data of *Oryza sativa* from NCBI GEO database (accession number GSE144419) [43] and CS endosperm RNA-seq from NCBI BioProject database (accession number PRJEB5135) [19]. Sequencing reads were cleaned with the fastp (version 0.20.0) [44] and Trim Galore (version 0.4.4), which eliminated bases with low quality scores (< 25) and irregular GC contents, sequencing adapters, and short reads. The remaining cleaned reads were mapped to the International Wheat Genome Sequencing Consortium (IWGSC) reference sequence (version 1.0) with the Burrows–Wheeler Aligner (version 0.7.17-r1188) [45] for the DAP-sequencing, ChIP-sequencing and DHS sequencing data. The HISAT2 program (version 2.2.1) [46] was used for mapping the RNA sequencing (RNA-seq) reads to the reference sequences. Reads with mapping quality less than 20 were removed. For ChIP-seq and DNase-seq and RNA-seq, the multi-mapped reads were directly removed.

The MACS (version 2.2.6) [47] program was used to identify the read-enriched regions (peaks) with the cutoff *P* < 1×10^−10^. For DAP-seq, the peaks detected from samples introduced with Halo tag only were considered as non-specific bindings, and TF peaks overlapping with peaks detected from Halo samples were removed for subsequent analysis. To quantify gene expression levels, the featureCount program of the Subread package (version 2.0.0) [48] was used to determine the RNA-seq read density for the genes. Integrative Genomics Viewer [49] was used to visualize the TFs binding, histone markers, gene expression and chromatin accessibility in the genome, and the number of reads at each position was normalized against the total number of reads (reads per million mapped reads).

### Processing of Hi-C data

Hi-C data of CS [50] (accession number GSE133885 deposited in NCBI GEO database) was processed as previously described [51]. Reads were aligned to the International Wheat Genome Sequencing Consortium reference sequence (version 1.0) and filtered by HiC-Pro (version 2.11.1) [52]. The default parameter “-q 10” was used to retain unique mapped read pairs. We used “findTADsAndLoops.pl” implemented in the Homer software to detect TAD-like domains [53]. We generated KR-normalized contact matrices with bin sizes set to 25 kb with Juicer and visualized the TADs with Juicerbox [54]. TAD-like domain boundaries were identified as 20 kb regions centered on the boundary points.

### Detection and enrichment analysis of transcription factor-binding motifs

The peaks were sorted by *q*-value and then by fold enrichment. The 600bp sequence centered on top 6000 peak summits were used to detect *de novo* motifs by MEME-ChIP [55] of the MEME software toolkit (version 5.1.1) and the enriched known motifs from JASPAR database were detected by AME [56] of the MEME software toolkit. The *de novo* motifs detected above were used to scan individual motif occurrences in genome with the FIMO program [57] of the MEME software toolkit. Motif logos were drawn by the R package motifStack (version 1.34.0) [58] and universalmotif (version 1.4.0).

### Calculation of the sequence conservation score

We completed a pair-wise comparison of the genome sequences with the NUCmer tool implemented in the MUMmer package [59]. For the diploid, tetraploid and hexaploid wheat comparation, the genome sequences from *T. urartu* (AA sub-genome, IGDBv1.0), *A. tauschii* (DD sub-genome, ASM34733 version 2), *T. turgidum* (AABB sub-genome, WEWSeq version 1.0) and *T. aestivum* (AABBDD sub-genome, IWGSC version 1.0) were used. The minimum sequence identity was set to 90 and each sub-genome was treated as an individual genome. Next, ROAST [60] was used to integrate pair-wise sequence alignments into a multiple sequence alignment. The multiple sequence alignment and tree data were fitted by PhyloFit, after which the conservation score was calculated with phastCons from the PHAST package [61].

### Detection of the subgenome homologous and nonhomologous regions

To detect subgenome homologous sequences, we used the subgenomic alignment results from NUCmer. Reciprocal aligned regions between subgenomes greater than 400 bp in length were defined as homologous regions. Other regions in the genome were defined as nonhomologous regions.

### Construction of co-expression network

We downloaded the hexaploid wheat expression data of 536 samples from Wheat Expression Browser (http://www.wheat-expression.com/) [18]. The samples and genes were filtered before constructing the network. Genes with TPM value less than 1 in at least 20 samples were removed and 200 samples were randomly selected to get a filtered expression matrix. Finally, 19,446 genes with high variance (top 25%) were retained. The WGCNA package (version 1.70.3) [62] was used to construct the co-expression network. An unsigned network was constructed by the blockwiseModules function, with the following parameters: power = 6; maxModuleSize = 6000; TOMType = “unsigned”, minModuleSize = 30; reassignThreshold = 0, mergeCutHeight = 0.25, numericLabels = TRUE; pamRespectsDendro = FALSE. If the co-expression partners of a gene could be defined by the abovementioned criteria, they would be assigned to the same module. Otherwise, the genes would be classified into module 0. All edges were ranked by the TOM value and top 80,000 edges were selected. The modules with HC and MC TFs (with 8,971 nodes) were visualized by Cytoscape (version 3.8.2) [63]. GO terms curated by GOMAP [64] were used to detect the overrepresented functional terms associated with the genes in each module.

### Sequence comparation of subgenome-nonhomologous TFBS

BLASTN was used to identify the similar nonhomologous TFBS pair within subgenome with the following parameters: E-value < 1e-30, identity > 70% and query coverage > 70%. The relationship of similar TFBS pairs (randomly selected 1,500 TFBS in each subgenome and *Arabidopsis thaliana*) were visualized by Circos [65].

To observe the expansion of TFBS, the multiple alignment of 500 randomly selected TFBS in each subgenome of each TF were performed by MAFFT (version v7.149b) [66] separately. The distance of each TFBS pair were calculated with distmat from EMBOSS (version 6.6.0.0) [67] with the Kimura 2-parameter correction. The sequence of randomly selected 500 TFBS of homologous TF in *Arabidopsis thaliana* and *Oryza sativa* were aligned and calculated the distance in the same way.

### Enrichment of specific TE families contributed to TF binding

TE annotation of CS was performed as previously described [51]. TE subfamilies account for more than 0.1% length of the genome were selected, and the enrichment scores (ES) between 98 TE subfamilies and 45 TFs were calculated. Enrichment of TE subfamilies was defined as:

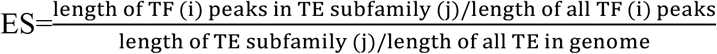

For TE enrichment analysis in the subgenome homologous and nonhomologous regions, merged TFBS of 45 TFs were used. For degenerated TE enrichment analysis, the non-degenerated TEs were used to calculate the length of TE subfamily (j) and the length of all TE in genome.

### Evolutionary analysis of enriched TE subfamilies

LTRharvest [68] was used to identify the full-length LTR of CS. Full-length RLC_famc1.4 were aligned with MAFFT separately. FastTree (version 2.1.10) was used to build the phylogenetic tree. The tree was visualized with R package ggtree (version 2.4.1) [69]. The insertion time was based on the divergence between the 5′ and 3′ LTRs and calculated with distmat from EMBOSS.

### Defination of subgenome regulatory divergence

Firstly, we calculated the regulatory activity of each TF to each target genes and then defined the subgenome regulatory divergence of each TF by comparing the regulatory activity of subgenome homologous genes.

The regulatory activity was quantified by TF affinity score (TFAS):

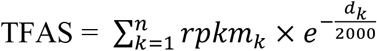

Where *d* is the distance of the peak summit to gene TSS and *rpkm* is the normalized reads count (reads per kilobase per million mapped reads) in peaks. The promoter was defined as 5 kb regions centered on the gene TSS. The peaks within promoter were considered as the peak with regulated effect of that gene. *n* is the count of total surrounding peaks. The TFAS of a gene was calculated by summarizing all *n* peaks. The TFAS of genes without TFBS in the promoter was 0.

Orthofinder [70] was used to identify the orthologous genes between subgenome. The orthogroups with only one copy in each subgenome(1:1:1) was defined as triads. The triads with all three genes TFAS less than 0.25 were filtered. We normalized the TFAS of genes in each triad by calculating the proportion of the TFAS of one subgenome among the three subgenomes. Subgenome balanced and unbalanced regulatory divergence pattern could be represented by seven standard TFAS proportion. The proportion [0.33, 0.33, 0.33] representing the balanced regulatory divergence pattern of subgenome (ABD). The proportion [0.5, 0.5, 0], [0.5, 0, 0.5], [0,0.5, 0.5] representing the unbalanced regulatory pattern-2 (AB, AD, BD) and [1, 0, 0], [0, 1, 0], [0, 0, 1] representing the unbalanced regulatory pattern-1 (A, B, D). The Euclidean distance from the normalized TFAS to seven standard coordinates were calculated for each triad. The subgenome regulatory divergence pattern was assigned to the pattern of standard TFAS proportion with the closest distance.

For regulatory divergence and expression divergence comparison, the regulatory divergence was quantified as the |log_2_ fold-change| of DAP read count in promoters between subgenome 1:1 orthologous genes. Orthologous pairs with TFAS greater than 0.25 of at least one gene were used. For each orthologous pair, we summarize the regulatory divergence of all targeting TFs. The expression divergence was quantified as the |log_2_ fold-change| of CS seedling between subgenome 1:1 orthologous genes.

### Definition of degenerated TE

For triad with unbalanced TE subfamily distribution in three subgenomes, BLASTN algorithm (version 2.9.0) was used to identify the degenerated TE by aligning the TE sequence of each TE subfamily in promoters of one or two subgenomes to promoters without that TE subfamily. The sequence of TE and degenerated TE in *URE* promoter were aligned by MAFFT and then visualized in Jalview (version 2.11.1.3) [71].

## Data access

The DAP-sequencing data generated in this study have been submitted to the NCBI Gene Expression Omnibus (GEO; https://www.ncbi.nlm.nih.gov/geo/) under accession number GSE192815 (https://www.ncbi.nlm.nih.gov/geo/query/acc.cgi?acc=GSE192815). Tracks for all sequencing data can be visualized through our local genome browser (http://bioinfo.sibs.ac.cn/dap-seq_CS_jbrowse/). histone ChIP-seq data of 7 typical tissues and 8 external stimuli, RNA-seq and DNase-seq data of Chinese Spring (CS) seedling are under accession numbers GSE139019 and GSE121903 in NCBI GEO database [11, 12]. Hi-C data of CS [50] is under accession number GSE133885 deposited in NCBI GEO database. The hexaploid wheat transcriptomic data of 536 samples were downloaded from Wheat Expression Browser (http://www.wheat-expression.com/) [18]. The *OsHOX24* ChIP-seq data of *Oryza sativa* is under accession number GSE144419 deposited in NCBI GEO database [43]. The TFBS of *Arabidopsis thaliana* were downloaded from Plant Cistrome Database (http://neomorph.salk.edu/dap_web/pages/index.php) [72].

## Acknowledgements

This study was supported by the National Science Fund for Excellent Young Scholars (32022012), National Natural Science Foundation of China (31921005), the Strategic Priority Research Program of the Chinese Academy of Sciences (XDA26030302 and XDB27010302). We thank Dr. Jizeng Jia from Chinese Academy of Agricultural Sciences for insightful comments. We thank Huang Tao for his help in maintaining the high performance computing server.

## Author contributions

Y.J.Z., Y.B.X. and Z.B.L. conceived and designed the experiments. W.L.Z, Y.P.T., Z.J.L., L.H.Y., Y.P., Y.L.Z., W.T. and Y.E.Z performed the experiments. Y.Y.Z., J.Y.L., Y.M., M.Y.W., H.S.D, Y.L.X., T.F.T., and Y.J.Z. analyzed the data. Y.J.Z. wrote the manuscript with input from all authors.

## Supplemental Figures for

**Supplemental Figure 1.**
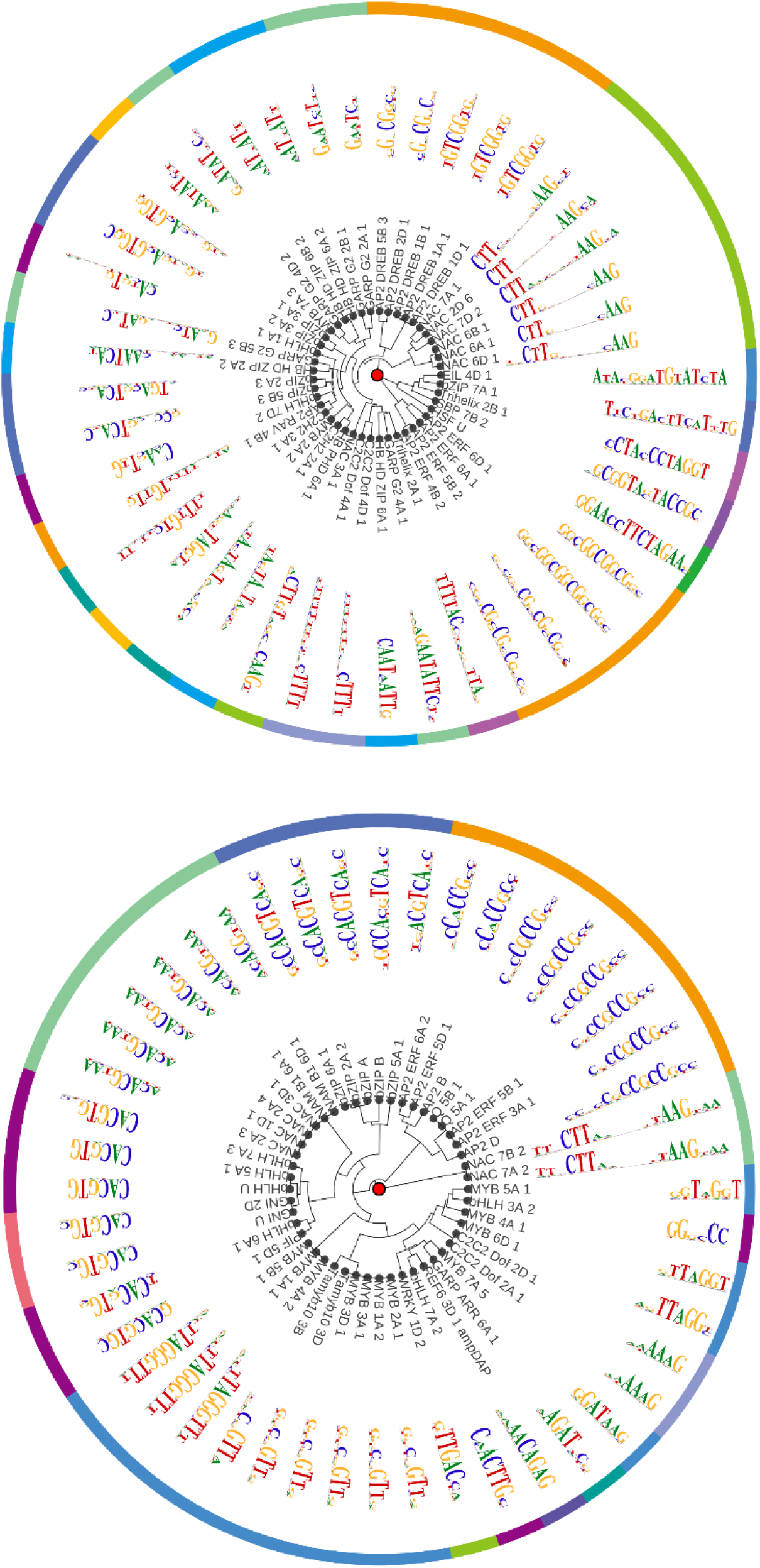
Clustering of the canonical motifs *de novo* identified in high confidence TFs peaks (top) or enriched in median confidence TF peaks (bottom).

**Supplemental Figure 2.**
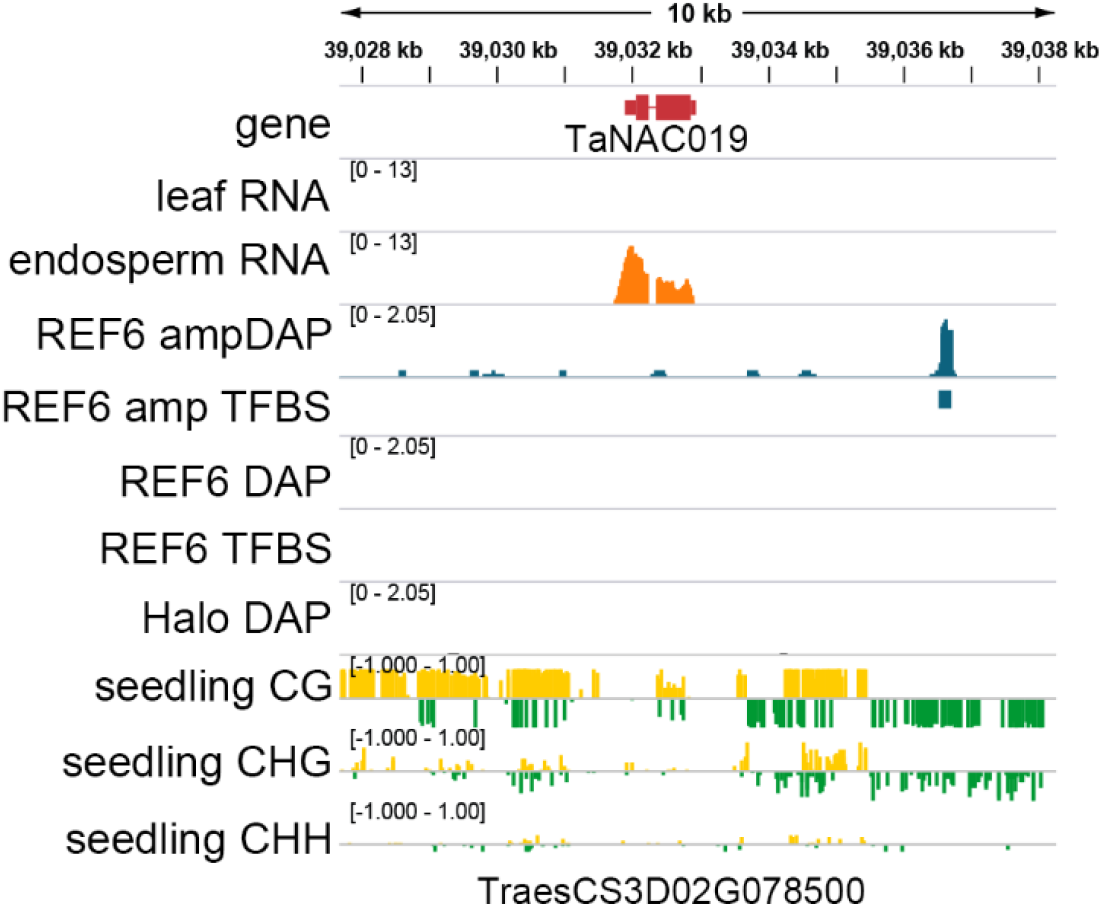
Genome tracks illustrating the low confidence (LC) TF REF6, whose binding is not enriched for the canonical motif of REF6 (CTCTGYTY), directly bind the promoter of endosperm specifically expressed gene *TaNAC019* when the DNA methylation of genome library was removed (REF6 ampDAP). Similarly, the hypo-methylated homologous gene *AT3G15170 (CUC1/ATNAC1)* gene body was also targeted by REF6 to activate cotyledon separation in Arabidopsis [1, 2].

**Supplemental Figure 3.**
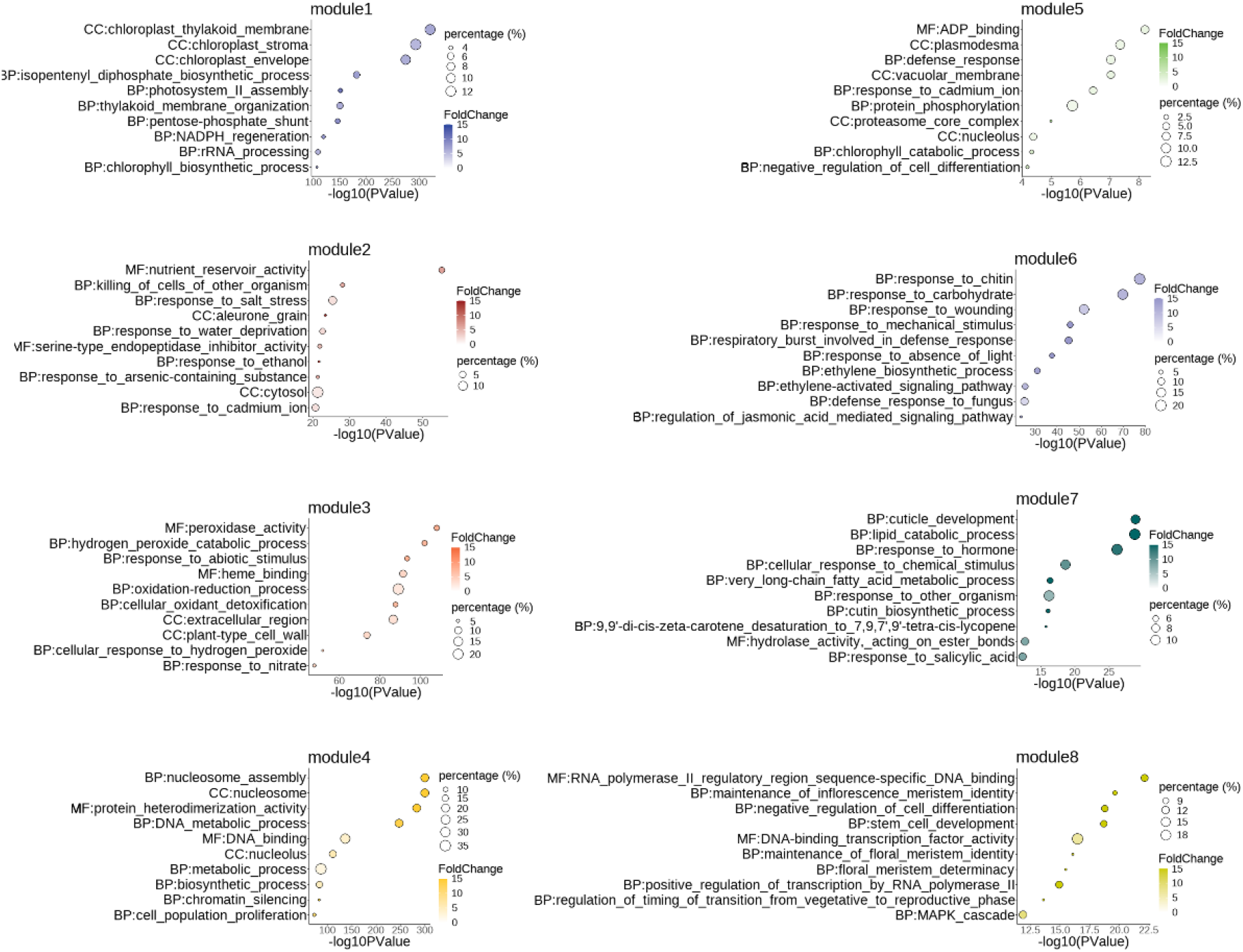
The top 10 enriched GO term ranked by *P* value of each co-expression network module.

**Supplemental Figure 4.**
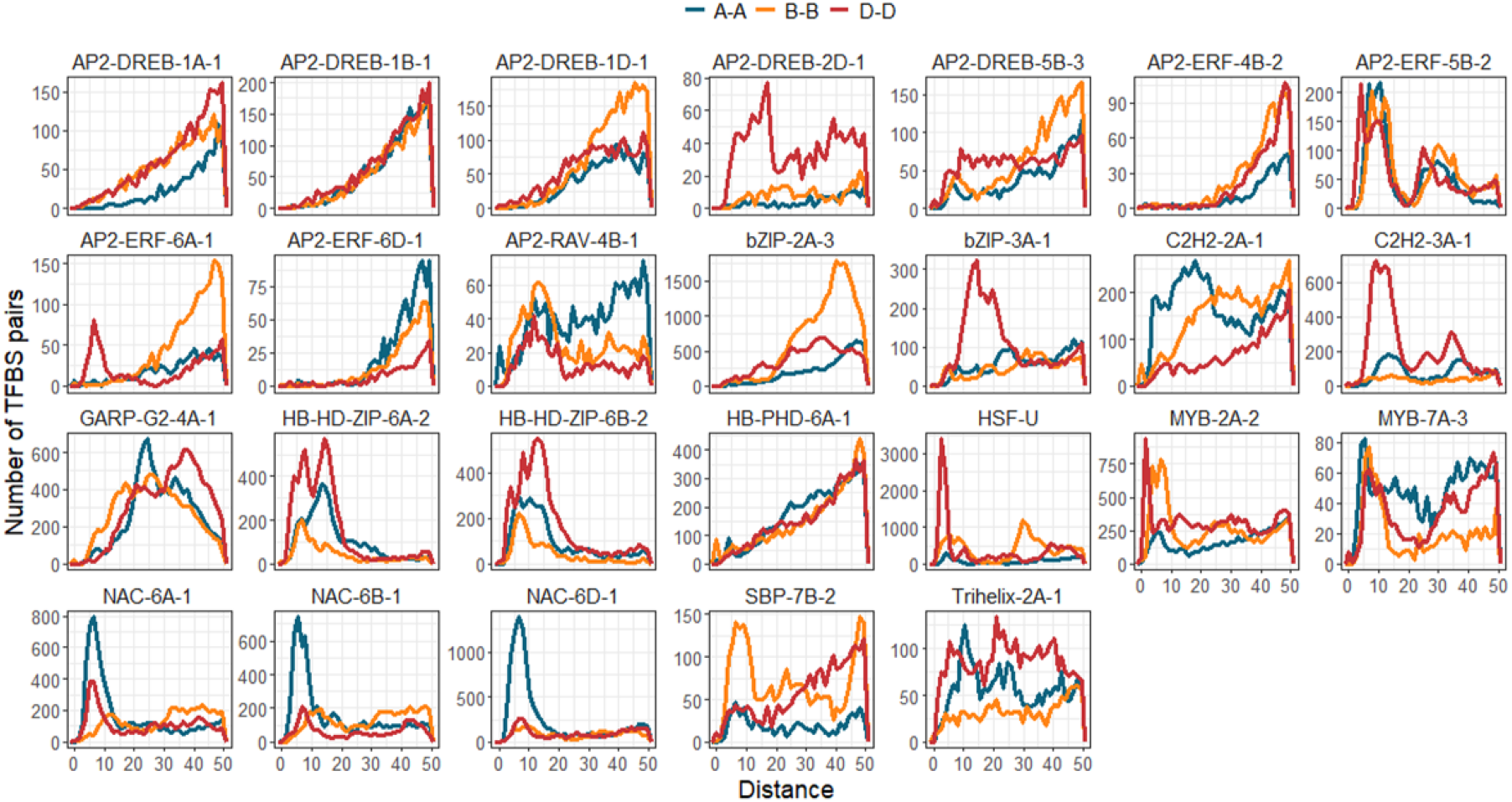
Sequence distance distribution between TFBS in each subgenome of CS.

**Supplemental Figure 5.**
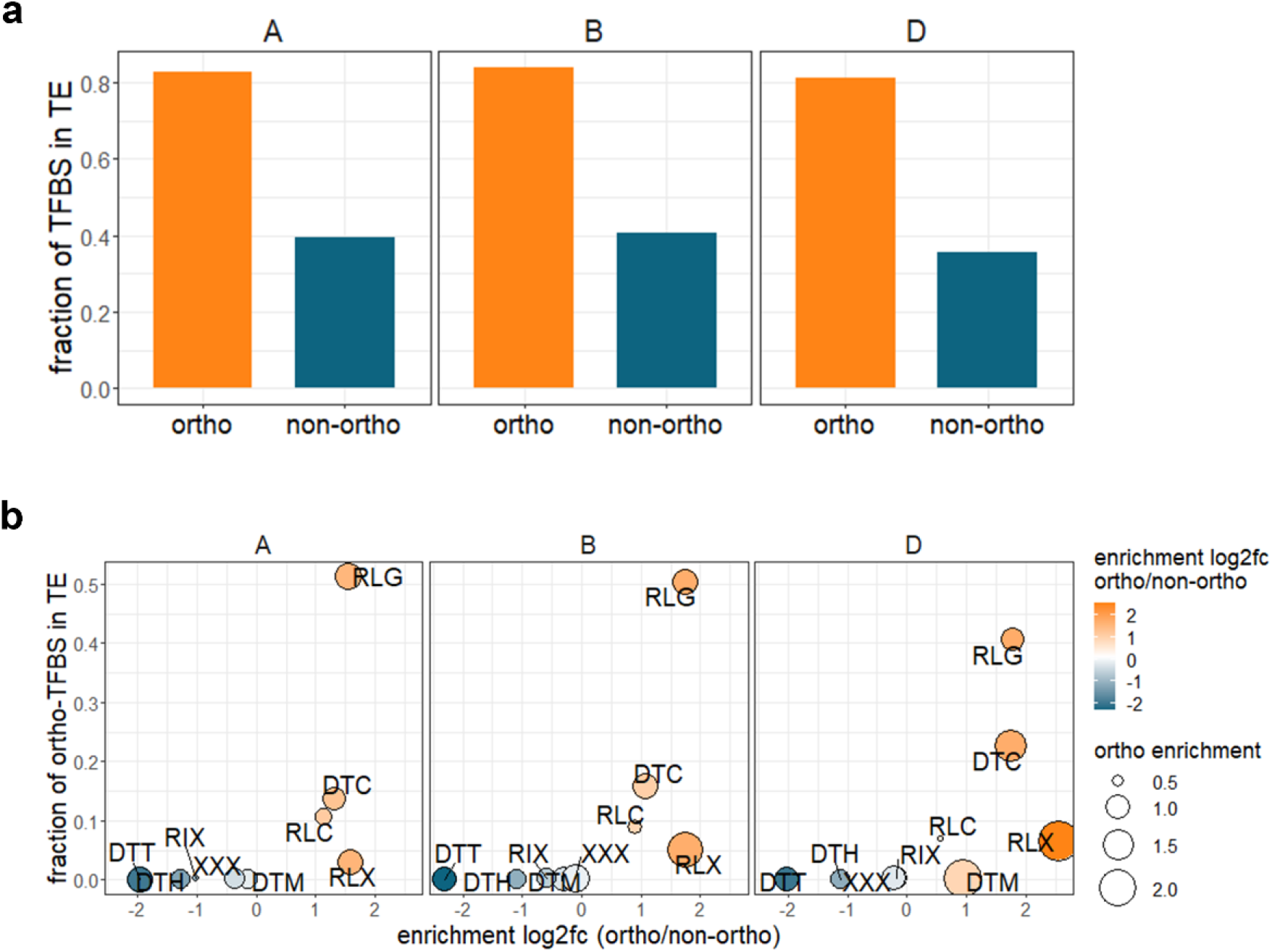
Fraction and enrichment of TE in ortho and non-ortho subgenome-nonhomologous TFBS. **a**, Fraction of expanded and non-expanded subgenome-nonhomologous TFBS in TE. Subgenome-nonhomologous TFBS were aligned within subgenome with BLASTN (see Methods). The TFBS pairs with similar sequence were identified as ortho TFBS and other TFBS were identified as non-ortho TFBS. **b**, Enrichment of TE families in ortho and non-ortho TFBS. The color bar and y-axis representing the log_2_ fold-change of enrichment score between ortho and non-ortho TFBS. The circle size representing the enrichment score of ortho TFBS. The x-axis representing the fraction of ortho TFBS in TE.

**Supplemental Figure 6.**
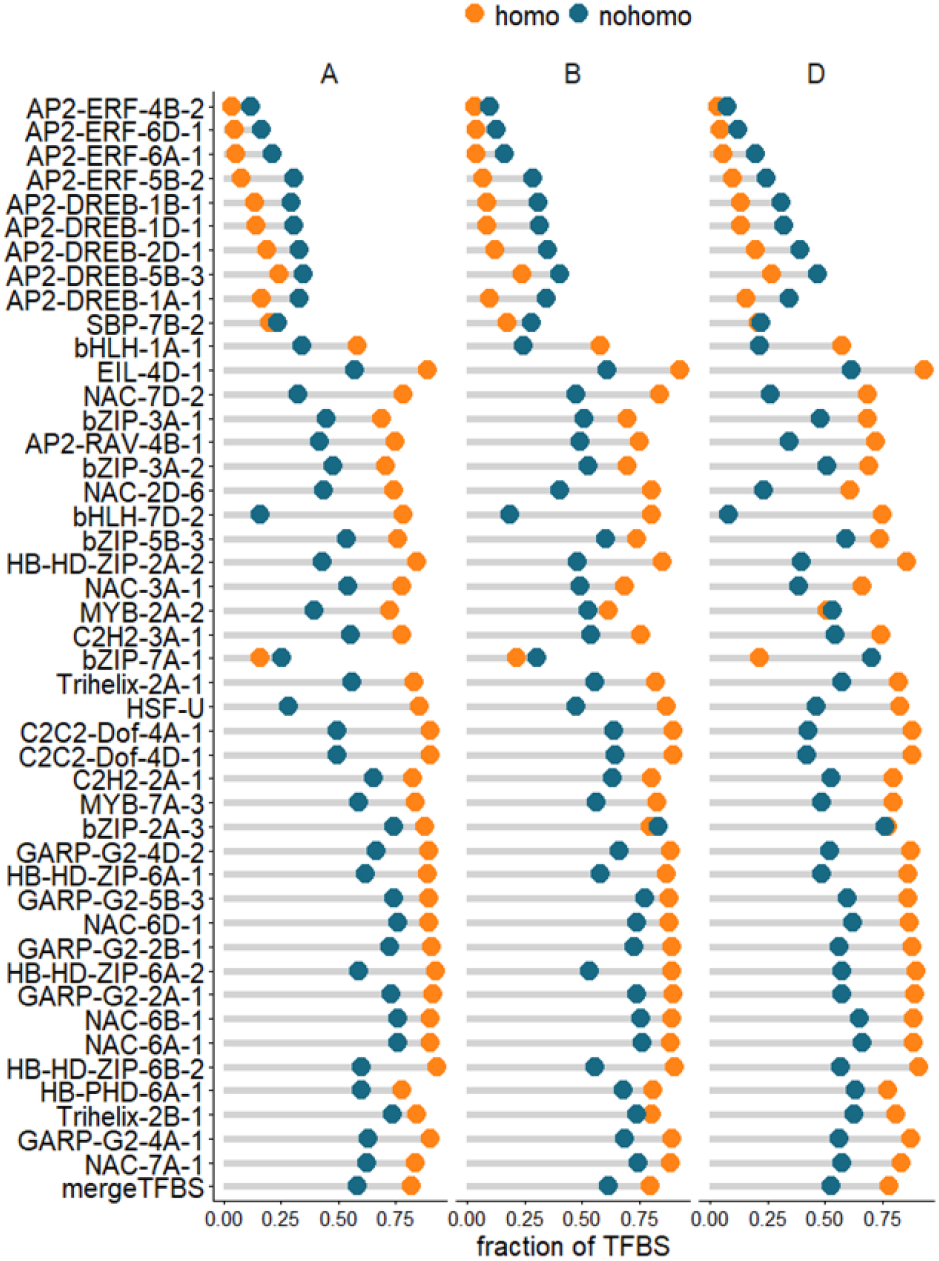
Fraction of subgenome-homologous and -nonhomologous TFBS embedded in TEs.

**Supplemental Figure 7.**
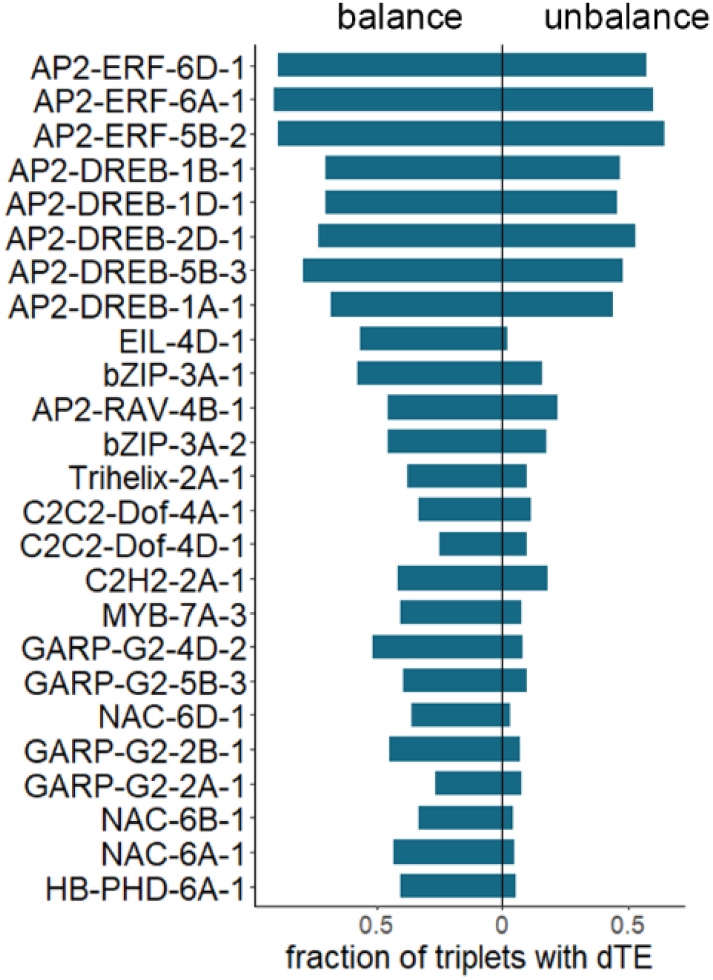
Fraction of targeted triplets with degenerated TE in balanced and unbalanced triplet.

**Supplemental Figure 8.**
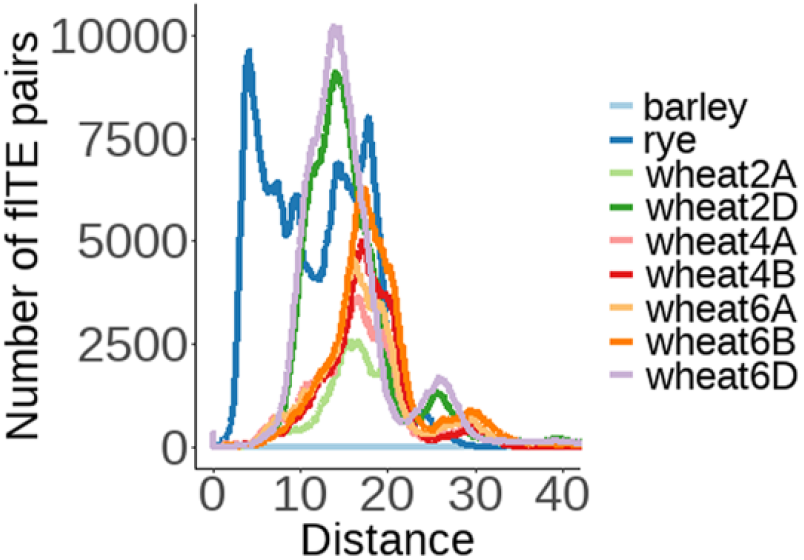
Frequency distribution of full-length TE sequence divergence of RLC_famc1.4 in Triticeae.

